# Uncovering thematic structure to link co-occurring taxa and predicted functional content in 16S rRNA marker gene surveys

**DOI:** 10.1101/146126

**Authors:** Stephen Woloszynek, Joshua Chang Mell, Gideon Simpson, Michael P. O’Connor, Gail L. Rosen

## Abstract

**Background:** Analysis of microbiome data involves identifying co-occurring groups of taxa associated with sample features of interest (*e.g.,* disease state). But elucidating key associations is often difficult since microbiome data are compositional, high dimensional, and sparse. Also, the configuration of co-occurring taxa may represent overlapping subcommunities that contribute to, for example, host status. Preserving the configuration of co-occurring microbes rather than detecting specific indicator species is more likely to facilitate biologically meaningful interpretations. In addition, analyses that utilize both taxonomic and predicted functional abundances typically independently characterize the taxonomic and functional profiles before linking them to sample information. This prevents investigators from identifying the specific functional components associate with which subsets of co-occurring taxa.

**Results:** We provide an approach to explore co-occurring taxa using “topics” generated via a topic model and then link these topics to specific sample classes (*e.g.,* diseased versus healthy). Rather than inferring predicted functional content independently from taxonomic abundances, we instead focus on inference of functional content within topics, which we parse by estimating pathway-topic interactions through a multilevel, fully Bayesian regression model. We apply our methods to two large publically available 16S amplicon sequencing datasets: an inflammatory bowel disease (IBD) dataset from Gevers *et al.* and data from the American Gut (AG) project. When applied to the Gevers *et al.* IBD study, we demonstrate that a topic highly associated with Crohn’s disease (CD) diagnosis is (1) dominated by a cluster of bacteria known to be linked with CD and (2) uniquely enriched for a subset of lipopolysaccharide (LPS) synthesis genes. In the AG data, our approach found that individuals with plant-based diets were enriched with Lachnospiraceae, *Roseburia*, *Blautia*, and *Ruminococcaceae*, as well as fluorobenzoate degradation pathways, whereas pathways involved in LPS biosynthesis were depleted.

**Conclusions:** We introduce an approach for uncovering latent thematic structure in the context of sample features for 16S rRNA surveys. Using our topic-model approach, investigators can (1) capture groups of co-occurring taxa termed topics, (2) uncover within-topic functional potential, and (3) identify gene sets that may guide future inquiry. These methods have been implemented in a freely available R package https://github.com/EESI/themetagenomics.

## BACKGROUND

With the decreasing cost of high-throughput sequencing, large datasets are becoming increasingly available, particularly microbiome datasets rich in sample data. These data include categorical and numeric features associated with each sample, which, in turn, may be linked to a set of taxonomic abundances that are derived from clustering sequencing reads. Typically, taxonomic marker genes, such as a portion of the 16S rRNA gene common to all bacteria, are used to perform the clustering based on a fixed degree of sequence similarity among reads. These clusters are termed Operational Taxonomic Units (OTUs), and each OTU corresponds to a taxonomic level, such as a genus.

Analysis of OTU abundances often involves identifying OTUs associated with specific sample features (*e.g.,* body site, disease presence, diet, age) via unsupervised exploratory methods such as principal component analysis, correspondence analysis, multidimensional scaling, and hierarchical clustering. Analyses may also include statistical inference strategies aimed at identifying differentially abundant OTUs and differences in alpha and beta diversity. Nevertheless, model building is hindered by the complexity inherent to abundance data of this type, which have a disproportionate number of OTUs relative to samples [1], a substantial degree of sparsity, and are typically strictly non-negative and constrained to sum to 1, *i.e.,* compositional [2, 3].

From an ecological perspective, co-occurring OTUs may represent related, overlapping groups of taxa that correlate with, for example, host status. Identifying important groups of co-occurring taxa, which we will refer to as “subcommunities,” facilitates a more biologically meaningful interpretation than identifying single indicator OTUs. This is because identifying subcommunities preserves the natural groupings of taxa when making inferences with respect to important sample features [4–7].

Still, suitable approaches are lacking that uncover the potential mechanistic relationships between subcommunities and sample features, namely how functional content within a subcommunity could correlate with sample features, such as genes belonging to a particular metabolic pathway correlating with disease status. That is, few methods successfully integrate subcommunity and sample features with functional profiles specific to these subcommunities.

In 16S rRNA survey approaches, analyses using taxonomic and functional abundance information typically involve independently inferring how the taxonomic and functional profiles of the samples associate with host features (Figure 1). The functional profiles are predicted based on taxonomic abundances with methods that use preexisting gene annotations. Examples of these methods include PICRUSt, Tax4fun, and Piphillin [8–10]. Because the inference of taxonomic and functional abundances occurs in two stages, it remains difficult to identify which functional content associates with which subsets of co-occurring taxa.

**Figure 1.**
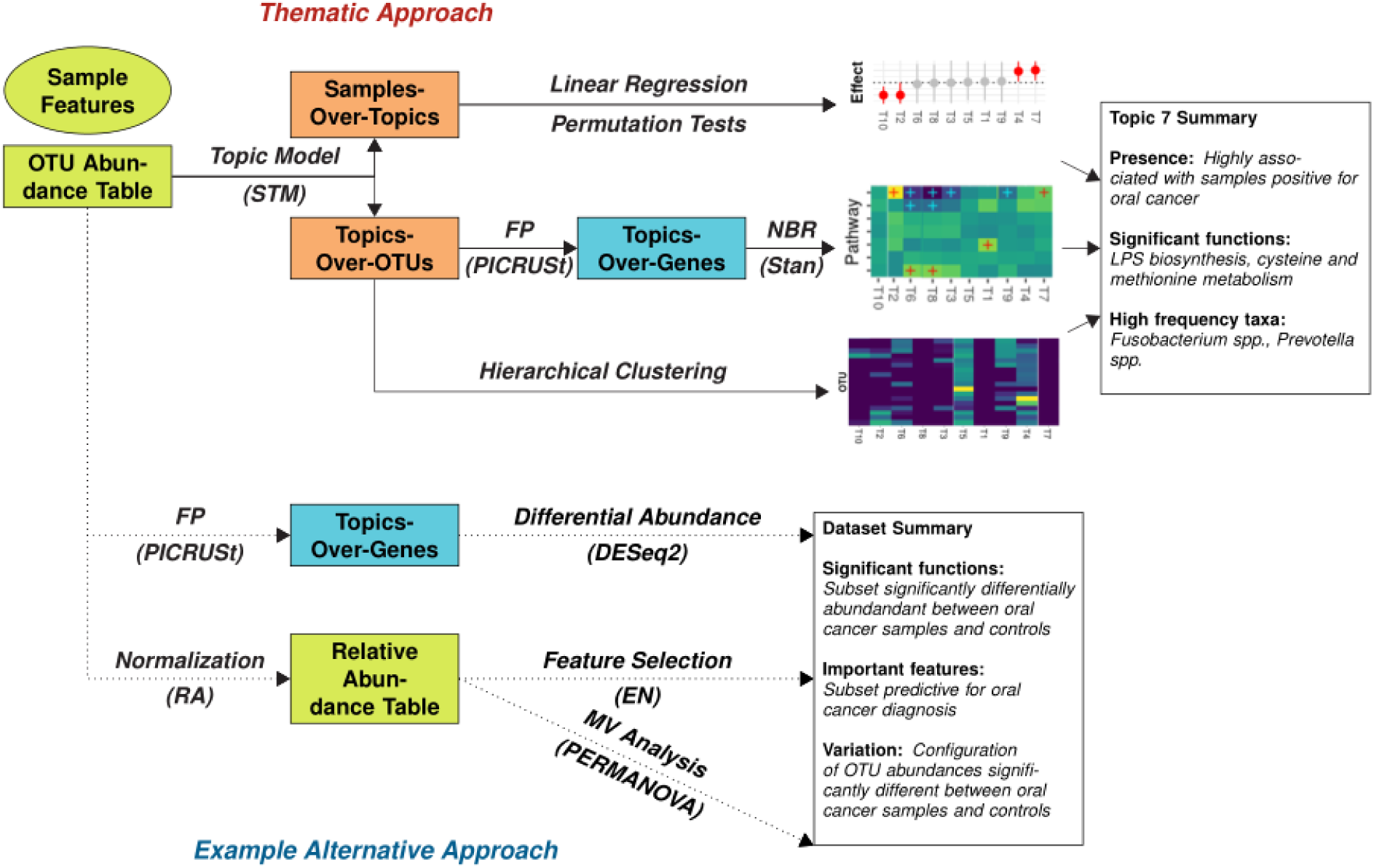
(Thematic Approach) Given a 16S rRNA abundance table, a topic model is used to uncover the thematic structure of the data in the form of two latent distributions: the samples-over-topics frequencies and the topics-over-OTUs frequencies. The samples-over-topics frequencies are regressed against sample features of interest to identify the strength of a topic-covariate relationship to rank topics (top). The topics-over-OTUs frequencies are used in a gene function prediction (FP) algorithm to predict gene content. Important functional categories are identified via a fully Bayesian multilevel negative binomial (NBR) regression model (middle). The topics-over-OTUs distribution is also hierarchically clustered to infer relationships between clusters of co-occurring OTUs and topics (bottom). The end result is the ability to identify key topics that associate clusters of bacteria, sample features of interest, and functional content. (Alternative Approach). An example alternative approach involves independently characterizing the taxonomic configuration and the predicted functional configuration of the OTU abundance table. Gene function prediction is performed on the full OTU abundance table, followed by a differential abundance analysis to infer differences in specific genes between sample features of interest (top). The OTU table is normalized to overcome library size inconsistencies and then analyzed via two methods: (1) an elastic net (EN) to find sparse sets of OTUs that are predictive for the sample feature of interest (middle) and (2) a multivariate (MV) analysis to identify relationships between beta diversity and the sample feature of interest (bottom). The end result are three analyses that summarize the data as a whole, unlike the thematic approach, which characterizes co-occurring sets of OTUs in three ways.

In the context of 16S rRNA surveys, we had two objectives: (1) implement a model framework that identifies subcommunities associated with specific sample features, and (2) uncover functional properties that further characterize these subcommunities.

To satisfy our first objective, we employed a topic model approach. Topic models have had considerable use in natural language processing, but have also shown promise as a method for exploring taxonomic abundance data. Knights *et al.* [11] used latent Dirichlet allocation (LDA) to infer the relative contributions of an unknown number of source environments to a set of indoor samples. Shafiei *et al.* [4], alternatively, took a supervised approach where they first trained their model on sets of co-occurring OTUs to learn how they correlate with sample classes of interest. They were then able to predict the class of new samples given the trained model.

Our approach uses a structural topic model (STM) [12], which generalizes previously described topic models such as LDA, the correlated topic model [13], and the Dirichlet-Multinomial regression topic model [14]. Like the Dirichlet-Multinomial regression topic model, the STM permits the use of sample features to help inform the frequency of topics occurring in a given sample. LDA, on the other hand, can only incorporate sample information if done in a two-stage process – first performing topic extraction, and then identifying linear relationships between the topic assignments and sample features [15]. A two-stage approach limits the breadth of sample information one can use, typically forcing the user to use only a single sample feature [12]. It also prevents propagating uncertainty throughout the model. Similar to the correlated topic model, the STM’s Logistic-Normal (LN) distribution defines the frequency of topics for a given sample and permits correlation between topics.

We use the STM to uncover a thematic representation of 16S rRNA survey abundance data and jointly measure its relationship with sample features (Figure 1). The aim is to cluster co-occurring OTUs into overlapping “topics,” where a given OTU can occur in multiple topics, albeit with varying frequency. The model also estimates the frequency of each topic in each sample based on that sample’s OTU abundances. Thus, the STM infers two latent, unobserved distributions from a table of OTU abundances: a samples-over-topics distribution (the frequency of each topic in each sample) and a topics-over-OTUs distribution (the frequency of each OTU in each topic). In addition, by utilizing sample features as covariates, the STM can also determine whether particular features increase or decrease the frequency of a topic occurring in a given sample, providing a means to identify topics associated with sample features.

Our second objective is to exploit the estimated topics-over-OTUs distribution, which dictates the taxonomic composition of each topic and therefore should capture meaningful subcommunities. We can therefore infer the functional content of individual topics using tools such as PICRUSt and a database of gene annotations. By identifying topics of interest based on their relationship to specific sample features and then inferring within-topic functional profiles, we can infer the specific functional content within subcommunities associated with sample features.

We apply our approach on two large 16S rRNA survey datasets: an inflammatory bowel disease (IBD) dataset from Gevers *et al.* [16] and data from the American Gut (AG) project. After confirming the generalizability of extracted topics, we identified distinct taxonomic subcommunities that, in the case of the Gevers *et al.* dataset, were consistent with published results. These subcommunities were composed of distinct functional profiles. Also, our approach provided gene sets specific to topics of interest that may warrant further exploration.

These methods have been implemented in a freely available R package themetagenomics: https://github.com/EESI/themetagenomics. In a companion paper, we performed simulations to further validate using topic models for 16S rRNA survey data and to determine a suitable normalization strategy (Woloszynek, S., Zhao, Z., Simpson, G., Mell, J. C., and Rosen, *in prep*).

## METHODS

### Review of the Structural Topic Model

The STM is a Bayesian generative model such that, given a set of *M* samples, each consisting of *N* OTUs, belonging to a vocabulary of *V* unique OTU IDs, *K* (chosen *a priori*) latent topics are assumed to be generated from the data. These topics consist of overlapping groups of co-occurring OTUs. A LN prior is placed on the samples-over-topics distribution. This allows for estimation of topic-topic correlations, giving a means to infer co-occurring topics across samples. The topics-over-OTUs distribution estimates deviation of OTU frequencies from the background OTU distribution [17]. Sparsity inducing priors are placed on the topics-over-OTUs distribution. This ensures a sparse set of estimates, ideal for high dimensional data. Lastly, word and topic assignments are both generated via multinomial distributions with *V* and *K* classes, respectively. For the relationships between topic model nomenclature and our terminology, see Table 1.

**Table 1.**
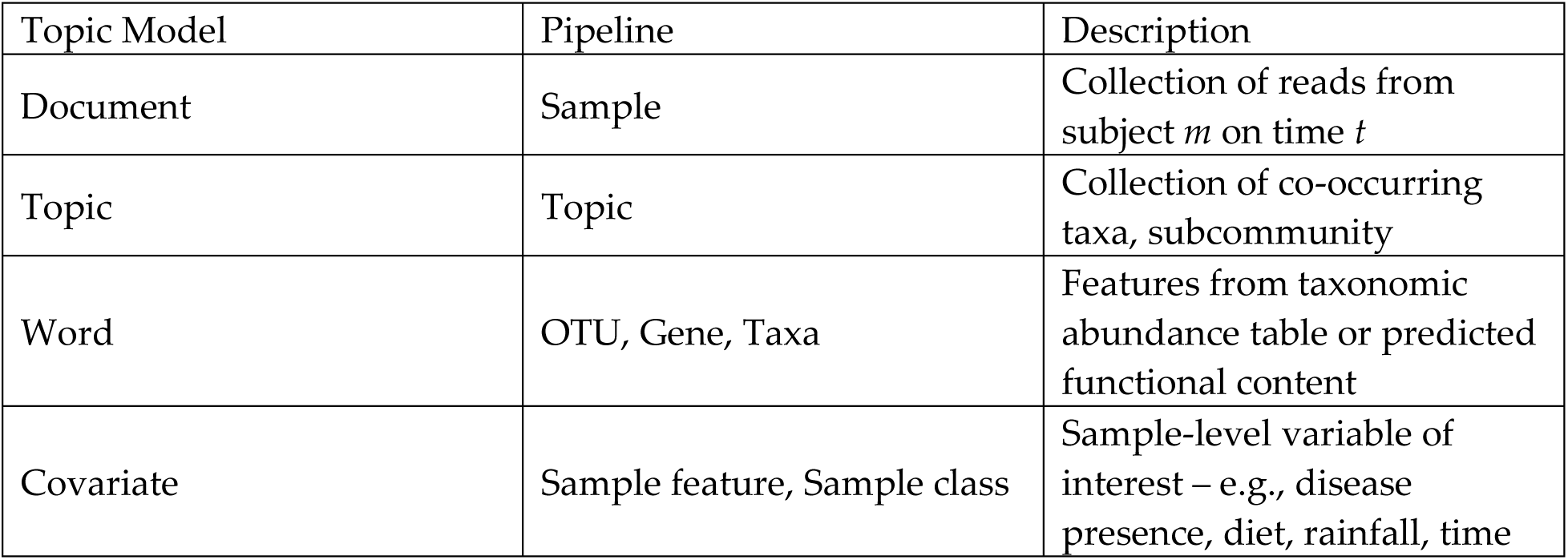
Relationship of Terms

| Topic Model | Pipeline | Description |
| --- | --- | --- |
| Document | Sample | Collection of reads from subject $m$ on time $t$ |
| Topic | Topic | Collection of co-occurring taxa, subcommunity |
| Word | OTU, Gene, Taxa | Features from taxonomic abundance table or predicted functional content |
| Covariate | Sample feature, Sample class | Sample-level variable of interest – e.g., disease presence, diet, rainfall, time |

The STM is estimated by a semi-collapsed variational expectation maximization procedure [18]. Convergence is reached when the relative change in the variational objective (*i.e.,* the estimated lower bound) in successive iterations falls below a predetermined tolerance.

### Datasets and Preprocessing

16S rRNA survey data from two human microbiome studies were downloaded from their corresponding repositories. The Gevers *et al.* dataset (“Gevers”) (PRJNA237362, 03/30/2016) is a multicohort, IBD dataset that includes control, Crohn’s disease (CD), and ulcerative colitis samples taken from multiple locations throughout the gastrointestinal tract [16]. The American Gut project (“AG”) (ERP012803, 02/21/2017) is a crowd-sourced dataset that includes user-submitted microbiome samples from a variety of body sites and self-reported subject information (http://americangut.org/).

#### Human gut microbiota from an inflammatory bowel disease cohort (Gevers)

Paired-end reads were joined and quality filtered (maximum unacceptable Phred quality score = 32; maximum number of consecutive low quality base calls before read truncation = 3; minimum number of consecutive high quality base calls included per read as a fraction of input read length = 0.75) using QIIME version 1.9.1. Closed-reference OTU picking was performed using SortMeRNA against GreenGenes v13.5 at 97% sequence identity. This was followed by copy number normalization via PICRUSt version 1.0.0 [19].

We selected only terminal ileum samples. Samples with fewer than 1000 total reads were omitted. We removed OTUs with fewer than 10 total reads across samples and OTUs that lacked a known classification at the phylum level.

#### Human gut microbiota from vegetarian and omnivore subjects (AG)

Quality trimming and filtering were performed in the following manner on single-end reads using the fastqFilter command found in the dada2 R package [20]. The first 10 bases were trimmed from each read. Reads were then trimmed to position 135 based on visualizing the quality score of sampled reads as a function of base position. Further truncation occurred at positions with quality scores less than or equal to 2. Any truncated read with total expected errors greater than 2 were removed. A portion of AG samples were affected by bacterial blooming during shipment. These reads were removed using the protocol provided in the AG documentation (02- filter_sequences_for_blooms.md).

OTU picking and copy number normalization were implemented as above. Samples with fewer than 1000 reads, and OTUs with fewer than 10 total reads across samples and lacking any known classification at the phylum level were discarded. We filtered samples falling into the “baby” age category (therefore the minimum age was 3) and retained only fecal samples. Within the diet category, unknown, vegetarian-with-shellfish, and omnivore-without-red-meat diets types were removed. We then merged vegan and vegetarian-without-shellfish into one class, resulting in a binary set of labels: “O” for omnivores and “V” for vegans and vegetarians.

### Fitting Structural Topic Models

Each resulting OTU table consisted of sets of raw counts normalized by 16S rRNA copy number. No other normalization was conducted based on the simulation results in Woloszynek *et al.* (*in prep*). A series of STMs with different parameterizations in terms of topic number (K ∊ 15, 25, 50, 75, 100, 150, 250) and sample features (*e.g.,* no features, indicators for presence of disease, diet type, etc.) were fit to the OTU tables. STMs were fit via the R package stm [21].

We evaluated each model fit for presence of overdispersed residuals. We also conducted permutation tests (permTest in the stm package) where the sample feature of interest is randomly assigned to a sample, prior to STM fitting. To compare parameterizations between models, we evaluated predictive performance using held-out likelihood estimation [15].

### Assessing Topic Generalizability

We performed classification to assess the generalizability of the extracted topics. No sample features were used as covariates. OTU abundances were split into 80/20 training-testing datasets. For different number of topics (K ∈ 15, 25, 50, 75, 100, 150), an STM was trained to estimate the topics-over-OTUs distribution. We then held this distribution fixed; hence, only the testing set’s samples-over-topics distribution was estimated. For both the training and testing sets, simulated posterior samples from the samples-over-topics distribution were averaged. The resulting posterior topic frequencies in the training set were then used as features to classify sample labels, similar to using 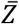 in supervised LDA [22]. Generalization (testing) error was assessed using the optimal parametrization based on cross-validation performance on the test set topic frequencies. Classification was performed using a random forest.

For the random forest, parameter tuning to determine the number of variables for each split was accomplished through repeated (10x) 10-fold cross-validation, using up- or down-sampling to overcome class imbalance (for Gevers and AG, respectively). We performed a parameter sweep over the number of randomly selected OTU features, while setting the number of trees fixed at 128. The optimal parameterizations were selected based on maximizing ROC area under the curve.

The performance of the STMs was compared to the performance using OTUs as features from the starting OTU abundance table. Separately, training and testing set OTU abundances were converted to relative abundances with the following equation: *OUT_n, m_*/∑_*n*_ *OUT_n, m_*. In words,

OTU *n* for sample *m* is scaled by the library size of sample *m* (the total abundance of sample *m*). The resulting OTU relative abundance tables were separately z-score normalized. Training cross-validation and testing using a random forest was then performed as above.

### Assessing Concentration of OTUs as a Function of Topic Number

For each STM (K ∈ 15, 25, 50, 75, 100, 150, 250), Shannon entropy was calculated for each topic in the topics-over-OTUs distribution. To compare mean entropy across STMs, we performed an ANOVA, followed by Tukey HSD post-hoc analysis.

### Identifying Within-Topic Clusters of High Frequency OTUs

Using the topics-over-OTUs distribution, we performed hierarchical clustering via Ward’s method on Bray-Curtis distances. We refer to high frequency groups of OTUs as “clusters.”

### Inferring Within-Topic Functional Potential

We obtained the topics-over-OTUs distribution for each fitted model and mapped the within-topic OTU probabilities to integers (“pseudo-counts”) using a constant: 10000 × *β*. A large constant was chosen to prevent low frequency OTUs from being set to zero, although their contribution to downstream analysis was likely negligible. Gene prediction was performed on each topic-OTU pseudo-count table using PICRUSt version 1.0.0 [8]. (Normalization of 16S copy number was performed prior to topic model fitting using PICRUSt.) Predicted gene content was classified in terms of KEGG orthology (KOs) [23].

### Identifying Topics of Interest

Topics of interest were identified using the samples-over-topics distribution as a matrix of *K* covariates in a linear regression model. Each column in the samples-over-topics distribution represents the frequency of topic *k* in sample *m*. The dependent variable chosen depended on the sample feature used as a covariate during model fitting. These include CD presence and PCDAI for Gevers and diet type for AG. We calculated 95% uncertainty intervals using an approximation that accounts for uncertainty in estimation of both the sample covariate coefficients and the topic frequencies. We will refer to these coefficients as “topic-effects.” Coefficients whose 95% uncertainty intervals do not span 0 will be referred to as “high ranking topics.”

### Identifying Functional Content that Distinguishes Topics

To determine which predicted functional gene content best distinguished topics, we used the following multilevel negative binomial regression model:

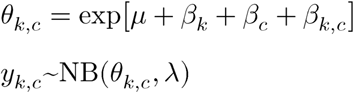

where *μ* is the intercept, *β_k_* is the per topic weight, *β_c_* is the per level-3 gene category weight, *β_k,c_* is the interaction weight for a given topic-function (gene category) combination, *y_k,c_* is the count for a given topic-function combination, and *λ* is the dispersion parameter. The intercept *μ* was given a Normal(0, 10) prior; all weights were given Normal 0, 2.5 priors; and the dispersion parameter *λ* was given a Cauchy(0, 5) prior.

Model inference was performed using Hamiltonian Monte Carlo in the R package rstanarm [24]. Convergence was evaluated across four parallel chains using diagnostic plots to assess mixing and by evaluating the Gelman-Rubin convergence diagnostic [25]. To reduce model size, we used genes belonging to only 15 (arbitrary number) level-2 KEGG pathway categories (Table S1). For large topic models, we fit only the top 25 topics, ranked in terms of topic-effects that measure the degree of association between sample-over-topic probabilities and our sample feature of interest.

### Assessing Relationships Between Sample Features of Interest and Taxonomic Abundance

To quantify the relationship between taxonomic abundance and continuous sample features (such as the Pediatric Crohn’s Disease Activity Index (PCDAI), a clinical measure of CD severity), we performed negative binomial regression (log-link), using sample library size (sum of OTU abundances across samples) as an offset. The family-wise error rate was adjusted via Bonferroni correction. Critical values for hypothesis testing was set at 0.05.

### Comparing Topic Taxonomic Profiles to a Network Approach

To further validate the clusters of high frequency taxa identified in the topics-over-OTUs distribution, we compared our results to those generated from an OTU-OTU association network on the copy number normalized OTU abundances using SPIEC-EASI’s neighborhood selection method (lambda.min.ratio=.01, nlambda=20) [26].

### Comparing Within-Topic Functional Profiles to an OTU-Abundance-Based Approach

We compared the results from the hierarchical negative binomial model to a differential abundance approach. We performed predicted functional content using PICRUSt on copy number normalized OTU abundances. The resulting functional abundances were collapsed into level-3 KEGG pathways. Note that, for consistency, we again restricted our genes to the 15 level-2 KEGG pathways used previously. The resulting level-3 pathway abundances underwent DESeq2 differential abundance analysis, which uses negative binomial regression and variance stabilizing transformations to infer the difference log-fold change of OTU abundance [27, 28]. The resulting p-values were corrected via the Bonferroni method. Adjusted p-values below 0.1 were considered significant.

### Packages utilized

All analysis was done in R version 3.2.3. Topic models, random forests, and NB regression models were fit using stm [21], caret [29], and rstanarm [24], respectively. AG filtering was performed using DADA2 [20]. SPIEC-EASI was fit using the SPIEC-EASI package [26]. DESeq2 differential abundance analysis was conducted with phyloseq [30]. Shannon entropy was performed with vegan [31].

### Implementation

Our approach can be implemented with themetagenomics, an R package that provides a topic model framework for microbiome abundance data, as well as functional prediction for 16S rRNA survey data. Users can choose between our C++ implementation of PICRUSt or Tax4fun functional prediction; therefore, both GreeneGenes and Silva taxonomic annotations are acceptable. Inference of topic-function interactions can be accomplished by maximum likelihood or Hamiltonian Monte Carlo using Rstan, where users can choose between Student-t, Laplace, and Normal prior distributions. The resulting topics, topic-effects, and topic-function interactions can then be explored with a variety of interactive Shiny apps.

## RESULTS

Here we explored the use of our structural topic model (STM) approach on publically available datasets of gut and fecal microbiota, first using the IBD data from Gevers *et al.* [32], and second using the dietary data from AG. For each dataset, we show that the topics extracted from the STM generalize well to test data not initially seen by the model, suggesting that co-occurrence profiles identified by the STM are robust to overfitting. Then, we apply our complete pipeline, where we successfully link, within a given topic, functional content, taxonomic co-occurrence profiles, and sample features of interest.

### Thematic Structure of IBD-Associated Microbiota (Gevers)

#### Dimensionality reduction using topics facilitates classification of CD diagnosis and generalizes well to test data

We sought to assess (1) if topics were associated with positive CD diagnosis (CD+) and (2) whether those topics generalize to new data – that is, did they capture meaningful information inherent to the data while ignoring characteristics associated exclusively with the fitted data. Note that, for this analysis, STM fitting was completely unsupervised, so no sample features were used as covariates.

The 80/20 training/testing splits for terminal ilium samples from Gevers are shown in Table S2. We hypothesized that using topics would outperform the relative abundance of OTUs as features for classifying CD diagnosis, since the relative abundance-based features are sparser. Both dimensionality and sparsity are reduced when using topics, since the size of the feature space is decreased through dimensionality reduction. There was little difference between the two approaches during training cross-validation with at least 25 topics (Figure S1, Table S3). During testing, however, topics outperformed OTUs, particularly in F1 score (a measure of performance that considers both sensitivity and positive predictive value), with scores of 0.808 for OTUs and at least 0.821 for all models with 25 or more topics (Table S4).

The largest discrepancy in classification performance between OTUs and topics was the proportion of true negatives out of total negative classifications (negative predictive value). The OTU model correctly identified CD- subjects only half the time (0.517), whereas the worst performing topic model (K=15) performed slightly better (0.526), and topic models improved as the number of topics increased: 0.655 (K=25), 0.559 (50), 0.577 (75), 0.682 (100), and 0.643 (150) (Table S4).

The substantially higher proportion of false negatives with the OTU model was likely due to its reliance on few, relatively rare taxa. For example, the random forest importance scores indicated that OTU 319708 (Clostridiaceae family) was the fourth most important feature for distinguishing CD+ from CD-. It was more than twice as common in CD- training samples, and more than 10% of correctly classified CD- samples contained this feature. When OTU 319708 was present in CD+ samples, it may have led these samples to be misclassified. Approximately 10% of misclassified CD+ samples contained this feature, and some of these samples contained it at a greater proportion than other samples in the training set. A similar scenario can be seen for OTU 186723 (Ruminococcaceae family), which received the highest random forest importance score for classifying CD. It was most common in CD+ samples; hence, its absence in CD+ samples resulted in false negatives.

#### Concentration of high probability OTUs across topics begins to plateau at 75 topics

We fit STM to the OTU abundance data and aimed to uncover how specific OTUs concentrate within topics as a function of topic number K (15, 25, 50, 75, 100, 150). We measured concentration using Shannon entropy. We define high quality topics as topics that place high probability on only a few OTUs; thus, high quality topics will have low entropy. A topic that is characterized by a small subset of OTUs is (1) more interpretable as a subcommunity and (2) contains more easily detectable associations with host features of interest.

For each K, we calculated the Shannon entropy for each topic, showing decreased entropy with increased topic number. This was supported by a one-way ANOVA (p<0.0001, F5,409=8.327) and post-hoc pairwise comparisons testing using Tukey HSD (α=0.05) (Figure S2). Among pairwise combinations, we found that models with 75 or more topics did not have significantly different Shannon entropies, leading us to focus our attention on topic models with at most 75 topics.

#### CD diagnosis was associated with distinct thematic profiles and hence distinct subcommunity structure

We implemented our full pipeline using sample features as covariates. Here, we used a binary indicator for CD diagnosis. We identified topics-of-interest based on their “topic-effects” – the regression coefficients estimated when regressing the samples-over-topics distribution against CD presence. We also performed permutation tests to ensure that detected topic-effects were not spurious effects. For model K25, we performed 25 permutations and calculated the mean posterior regression coefficient for each topic. Of the 25 topics, the 95% uncertainty intervals for 8 topics did not span 0 (Figure S3). We consider these “high ranking topics.” Topics T15, T12, T2, and T14 had estimates greater than 0, whereas topics T11, T25, T13, and T19 had estimates less than 0. For K75, 14 topics did not span 0 (Figure S4).

The posterior predictive distribution of each sample’s topic assignment is shown in Figure 2 for STMs K25 and K75. Each sample is plotted based on its PCDAI, which is 0 for CD- samples and increases as CD severity increases. Topics are ordered in terms of their topic-effects. Both panels demonstrate that as PCDAI increases, the thematic profile changes towards topics with higher correlations to CD+. Also, high ranking topics (cyan and red points for CD- and CD+, respectively) concentrate at the extremes, suggesting that the OTU abundance profiles for healthy and high severity CD samples were more precisely captured by the STM. Notably, there was clearer separation between CD- and CD+ associated high ranking topics for the K25 model. The transition point can clearly be seen at approximately PCDAI=35. Based on this result, we will hence focus on the K25 model.

**Figure 2.**
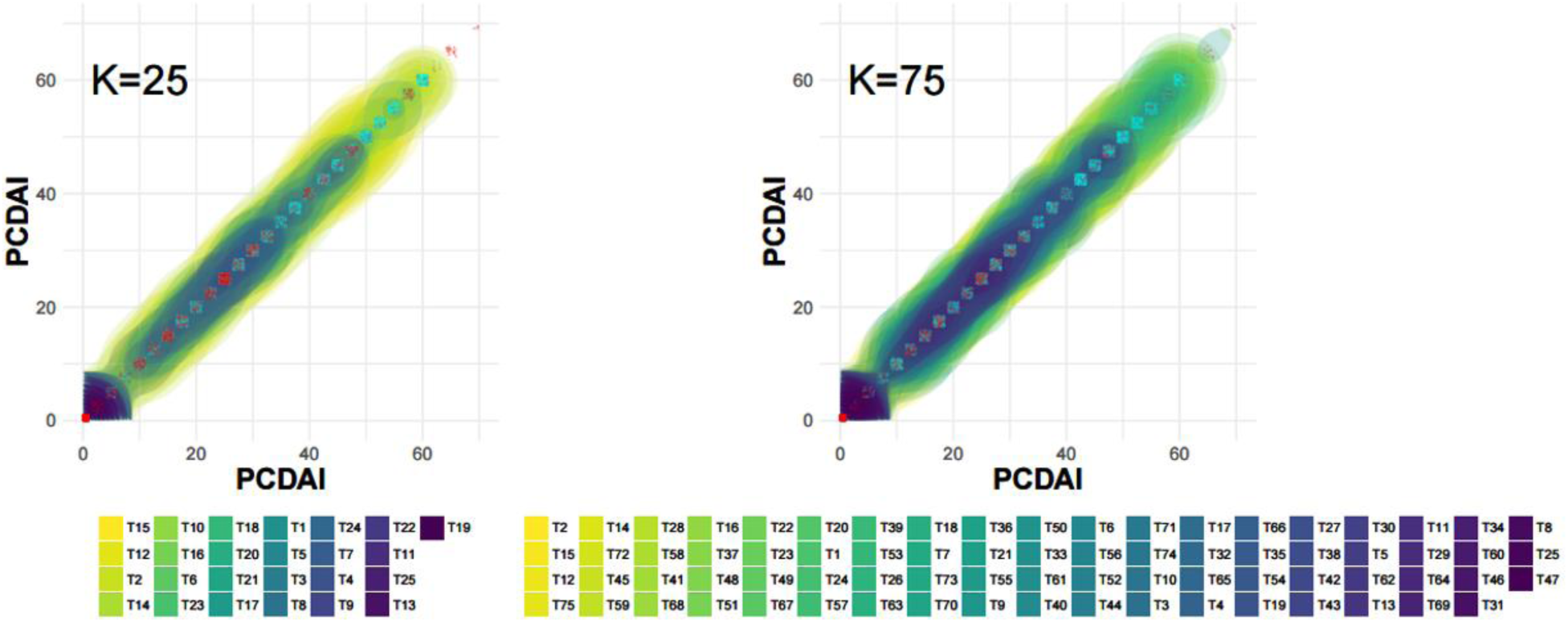
The posterior predictive distribution (PPD) of topic assignments for each sample in Gevers with reported PCDAI. Shown are the PPDs for K25 and K75. CD- samples were set to PCDAI=0. Topics are ordered based on their topic-effect, the regression weights estimated by regressing the samples-over-topics distribution against CD presence. Thus, left-most topics are most associated with CD- samples, whereas right-most topics are most associated with CD+ samples. The cyan (CD-) and red (CD+) points indicate if a topic, drawn from the PPD for a given PCDAI value, were high ranking topics (topics that did not span 0 at 95% uncertainty for permutation tests).

Focusing on the high ranking topics for K25 (T19, T13, T25, T11; T14, T2, T12, T15), we identified multiple clusters of bacterial species (via hierarchical clustering) that disproportionately dominated the high ranking topics associated with CD+ (Figure 3A). T2 contained a cluster dominated by *Enterobacteriaceae* taxa, whereas T12’s cluster contained a mixture of *Fusobacteria* and *Enterobacteriaceae*. The T15 cluster contained *Haemophilus* spp., *Neisseria*, *Fusobacteria*, and *Streptococcus*, all of which were noted as having a positive correlation with CD+ subjects in Gevers *et al.*, as well as *Aggregatibacter*, a genus reportedly associated with colorectal cancer [33].

**Figure 3A,B.**
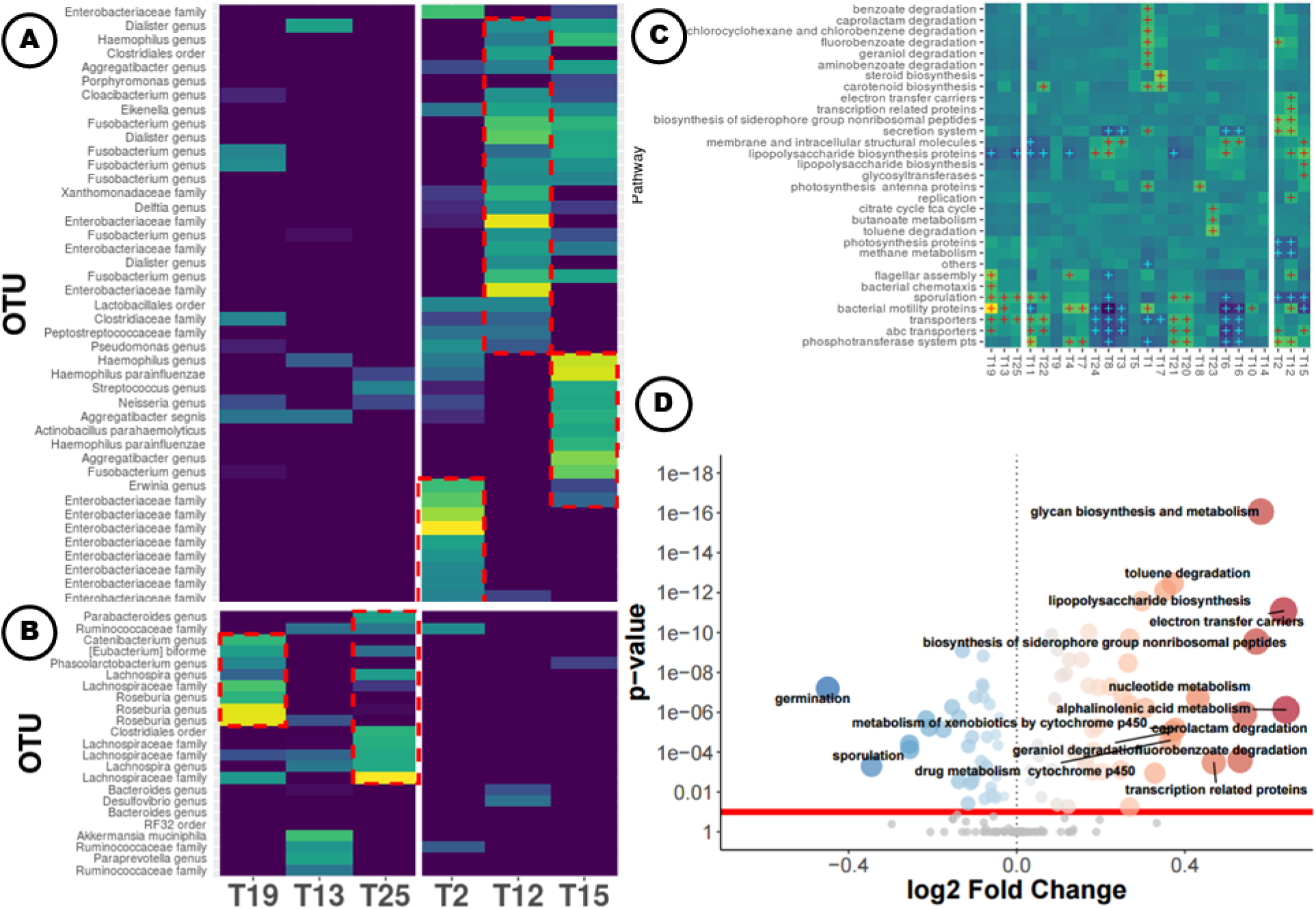
Subsections of the heatmap for the Gevers of the topics-over-OTUs distribution (K25) in log space. Shown are the top 3 topics associated with CD- and CD+, ordered by topic-effects (left to right, respectively, separated by the white line). Clusters of interest are marked with red dotted lines. Clustering was performed via Ward’s method on Bray-Curtis distances. Low probabilities (p < 1x10^-5^) are set to 0 to minimize the range of the color gradient to ease visualization. Yellow=high probability, Blue=low probability. Figure 3C. Level-3 pathway category-topic interaction regression coefficients from the multiple level negative binomial model. KEGG information was predicted via PICRUSt on the topics-over-OTUs distribution. Clustering was performed via Ward’s method on Bray-Curtis distances. Red and blue crosses indicate estimated pathway-topic interaction weights that do not span 0 at 80% uncertainty and are positive or negative, respectively. Only pathways with at least one such combination are shown. Yellow=large positive weight estimate, Blue=large negative weight estimate. Figure 3D. Volcano plot showing DESeq2 results for differentially abundant predicted level-3 KEGG categories. Functions were predicted using PICRUSt on the copy number normalized OTU abundance table. Blue and red points represent categories significantly enriched for CD- and CD+, respectively. Gray points are categories with p-values greater than 0.1 after Bonferroni correction.

Given that T15 contains a cluster of bacteria known for their association with bowel inflammation and this topic occurs disproportionately in subjects with greater disease severity, we asked whether the abundances of the OTUs in T15 correlated with PCDAI. After performing negative binomial regression (Figure 4), we identified significant positive trends as a function of PCDAI for *Aggregatibacter* (p<0.0001, β=0.089, Z=5.285), *Erwinia* (p=0.0004, β=0.103, Z=4.116), *Fusobacterium* (p=0.0001, β=0.081, Z=6.354), and *Haemophilus* (p=0.0484, β=0.0264, Z=2.847).

**Figure 4.**
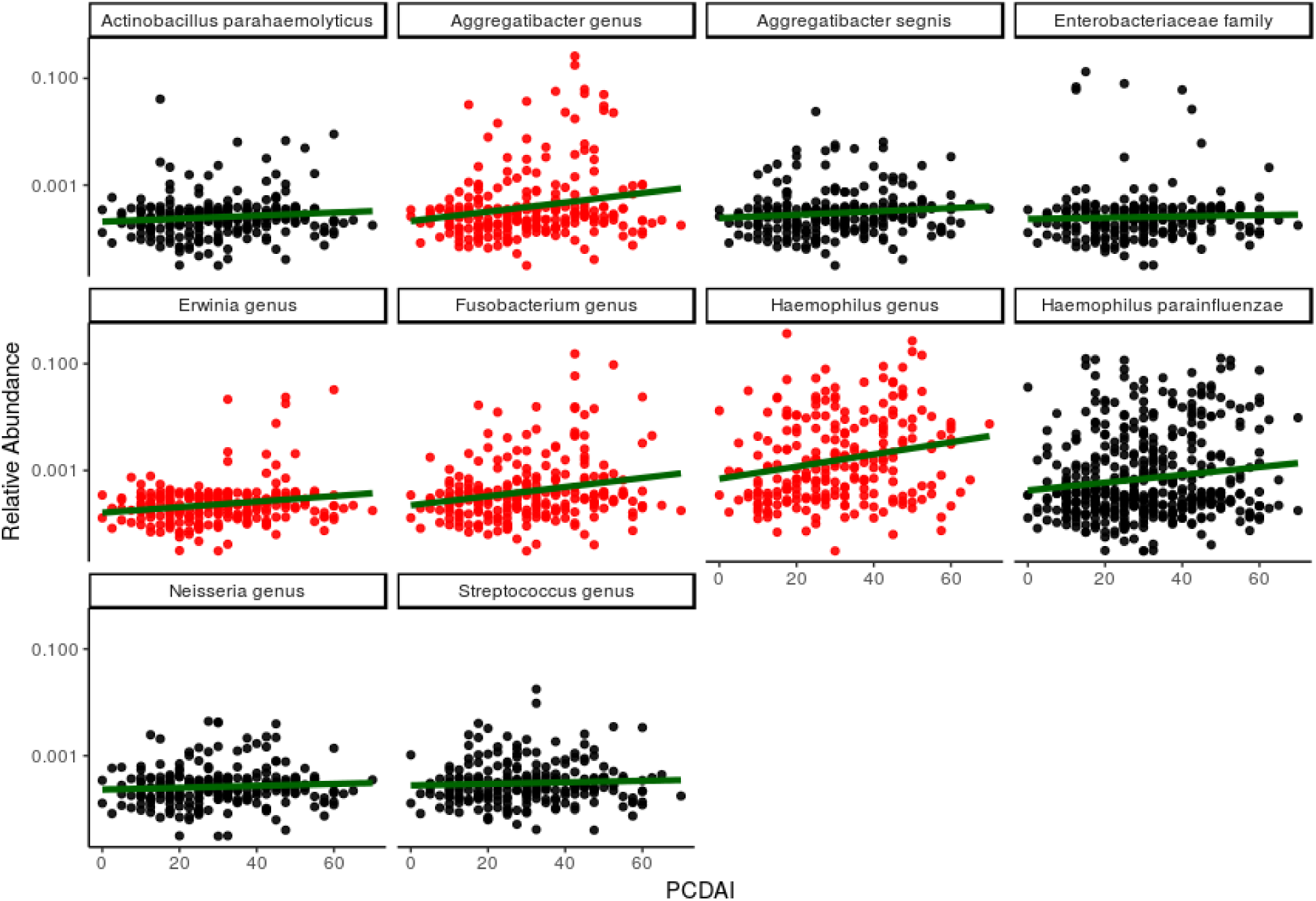
Scatterplots of Gevers data for the relative abundance of taxa that compose a high probability cluster in T15 versus PCDAI, a clinical measure of CD disease burden. Red points reflect significance (alpha=.05) for negative binomial regression (log linked, sample coverage offset) with Bonferroni correction.

The high ranking topics for CD-, on the other hand, were dominated by taxa belonging to *Lachnospiraceae*, *Roseburia*, *Rubinococcus*, *Blautia*, *Bacteroidetes*, and *Coprococcus*, all of which were noted by Gevers *et al.* as being negativity associated with CD (Figure 2B). In addition to these taxa, *Akkermania*, *Dialister*, and *Dorea* contributed to these topics, which is consistent with the findings of Lewis *et al.* who found a reduction of these taxa in CD+ subjects [34].

#### Within-topic co-occurrence profiles were consistent with SPIEC-EASI

We compared topics to the correlations obtained via a network approach. The edges in the SPIEC-EASI network for the clusters of high probability OTUs in our high ranking topics are shown in Figure S5. For each of these topic clusters, the majority of taxa were connected by a non-zero edge (Table S5). Of the 11 taxa in the T15 cluster, 8 had first-order connections (direct connections to other taxa within the cluster, OTU_c_-OTU_c’_), whereas 9 had second-order connections (indirect connections to other taxa within the cluster via an intermediate OTU not present in the cluster, OTUc-OTUnc-OTU_c’_). The two OTUs connected by the largest edge weight, *H. parainfluenzae* and *Haemophilus spp.*, had the highest frequencies in T15, 0.320 and 0.245, respectively. Of topics T15, T12, T2, T19, T13, and T25, none had more than one OTU with zero connections or fewer than 75% of taxa joined by first-order connections. The taxa that lacked within-cluster connections generally had low topic frequencies, with one exception, *Catenibacterium spp.* in T19. Taken together, this reaffirms that the within-topic co-occurrence profiles are consistent with alternative approaches.

#### Predicted functional potential of notable topics further elucidated their association with CD

We sought to further explore the functional content of topics, thereby exploiting the posterior estimates of the STM in a way other approaches have not. To do so, we predicted the functional content within topics using PICRUSt, which infers the metabolic potential of a microbial community by matching taxonomic classifications made from 16S rRNA gene sequences with a closely related reference genome annotation. We then performed a fully Bayesian multilevel regression analysis on the predicted abundances of each gene to identify topic-function interactions. This identified the bacterial taxa (irrespective of taxonomy) that drive the functional associations.

Like Gevers *et al.*, we identified an increase in membrane transport associated with CD+ subjects’ gut microbiome; however, through our approach, we were able to pinpoint the specific topics (*i.e.,* subcommunities) associated with the enrichment of these functional categories, topics T2 and T12 (Figure 2C). We then could link enrichment of membrane transport genes to the taxa that were also enriched in this topic. For example, topics T2 and T12 were dominated by Enterobacteriaceae. The Enterobacteriaceae-enriched topics (T2, T12) were also enriched for siderophore and secretion system related genes. Like T2 and T12, T15 was highly associated with CD+; however, it was less enriched for membrane transport genes. This suggests that the cluster of bacteria found in T15 (*Haemophilus* spp., *Neisseria*, and *Fusobacteria*) may have contributed less to the shift of transport genes reported by Gevers *et al.* and instead have distinct functional associations with CD.

The largest topic-function interaction effect was found in T19 for genes encoding bacterial motility proteins. For T19, three motility-related KEGG categories (bacterial motility proteins, bacterial chemotaxis, flagellar assembly) had topic-function interaction effects that did not span 0 at 80% uncertainty, suggesting that T19 was more enriched in cell motility genes relative to all other topics. The gene functions inferred for T19 are consistent with this taxonomic profile, consisting of motile bacteria belonging to Lachnospiraceae, Roseburia, and Clostridiales. Enrichment of two lipopolysaccharide (LPS) synthesis categories were associated with CD+ topics; however, one of these categories was specific for only T15 (Table S7).

#### More functional categories were significant via a DESeq2 approach on the OTU abundance table

We compared our within-topic functional profiles to the results obtained by performing PICRUSt on the copy-number normalized OTU abundance table and then performing a DESeq2 differential abundance analysis. Of the 160 level-3 KEGG categories, more than half (87) were found significant (α < 0.1), in the DESeq2 approach, leading to a difficulty in making meaning out of the data (Figure 3D). Pathways with the largest log-fold change (LFC) associated with CD+ samples included degradation pathways (caprolactam, LFC=0.542; fluorobenzoate, 0.532; geraniol, 0.371; and toluene degradation, 0.371), alphalinolenic acid metabolism (0.641), and electron transfer carriers (0.635).

Interestingly, the degradation pathways associated with CD+ also demonstrated strong topic-function interaction effects; however, they associated most strongly with T1, a topic unrelated to CD presence. Predicted electron transfer carrier genes were identified by both approaches, but the topic model approach isolated the effect to T12, placing high probability on bacteria that are also enriched for functions linked to secretion systems, LPS biosynthesis, and motility.

The topic-free DESeq2 approach also found fewer categories associated with CD- that had large LFC. For example, only one category had a LFC less than -0.4, whereas there were 8 greater than 0.-4 (enrichment in CD+). The categories with the largest LFCs relative to CD- included germination (LFC=-0.450) and sporulation (-0.346). Similarly, the topic model identified 10 topics with functional profiles significantly enriched or depleted in sporulation genes, three of which were associated with CD- samples. Multiple topics demonstrated an inverse relationship between sporulation and LPS genes, such that topics that contained taxa enriched in one were depleted in the other.

### Thematic Structure of Diet-Associated Microbiota (AG)

Despite consisting of far more samples, the AG dataset, split into omnivore (O) and vegetarian (V) diet groups from self-reported dietary information, offered a new challenge for our approach, given that there were far more data and taxonomic features (OTUs), as well as severe imbalance between classes. Of the 4864 samples that fit into our diet groups, 4527 were identified as O and only 337 as V samples. This disparity of group sizes limits the utility of comparing group means, particularly as an estimate of covariate effects [35].

#### BMI is a potential confounder

Before applying our pipeline, we considered potential sources of confounding. Male and female samples were distributed similarly with respect to diet (Table S9; Figure S6). There was no significant difference in mean age between diet groups (t=-0.03, df=373.93, p=0.98). Sample body mass index (BMI) was not normally distributed (Shapiro-Wilk: W=0.86, p<0.001) and was plagued with many mislabeled heights and weights (Figure S7). After attempting to remove samples we deemed unreliable (age < 17y, height < 1.4m, height > 2.2m, weight > 200kg), we still found a significant mean difference in BMI between diet groups via a Mann Whitney U test (p<0.001). Despite this difference in BMI, we removed no additional samples since we were concerned about further worsening the imbalance between diet groups.

#### Classification using topics is less conservative and, for low dimensional models, less generalizable

Unlike Gevers, models with fewer topics (K < 75) generalized poorly compared to using OTUs as features for classification, which may be due to AG having nearly 3-times as many unique OTUs, causing too few topics to dampen any meaningful signal (Table S11). Interestingly, all parameterizations outperformed the raw data in terms of sensitivity but not specificity (Table S11), suggesting that classification using OTU features is more conservative.

#### Diet was associated with specific taxonomic and functional profiles

We will report the results from a 100 topic STM, fit with the binary diet information as a covariate. As before, we identified our topics-of-interest by regressing the samples-over-topics distribution against diet and further validated these results via permutation tests, resulting in 9 high ranking topics, 5 of which were associated with the O group, and 4 with the V group (Figure S9).

Across the 9 topics, members of the family Lachnospiraceae were well represented, which is not surprising given that it typically accounts for over half of bacteria in healthy human fecal samples [36]. Within the topics, we identified roughly 11 clusters of interest that contained high frequency taxa.

The topic most associated with the V group T61, was dominated by taxa belonging to Lachnospiraceae, but was also enriched for *Roseburia*, *Blautia*, and *Ruminococcaceae* (Figure 5A). T61’s association with the V diet is consistent with literature linking *Roseburia* and *Ruminococcaceae* to starch and plant polysaccharide metabolism [37] and *Roseburia* and *Blautia* to whole grains [38, 39]. Also, consistent with this topic being dominated by Gram positive bacteria, we identified a significant depletion in LPS biosynthesis genes. (Figure 5B).

**Figure 5A.**
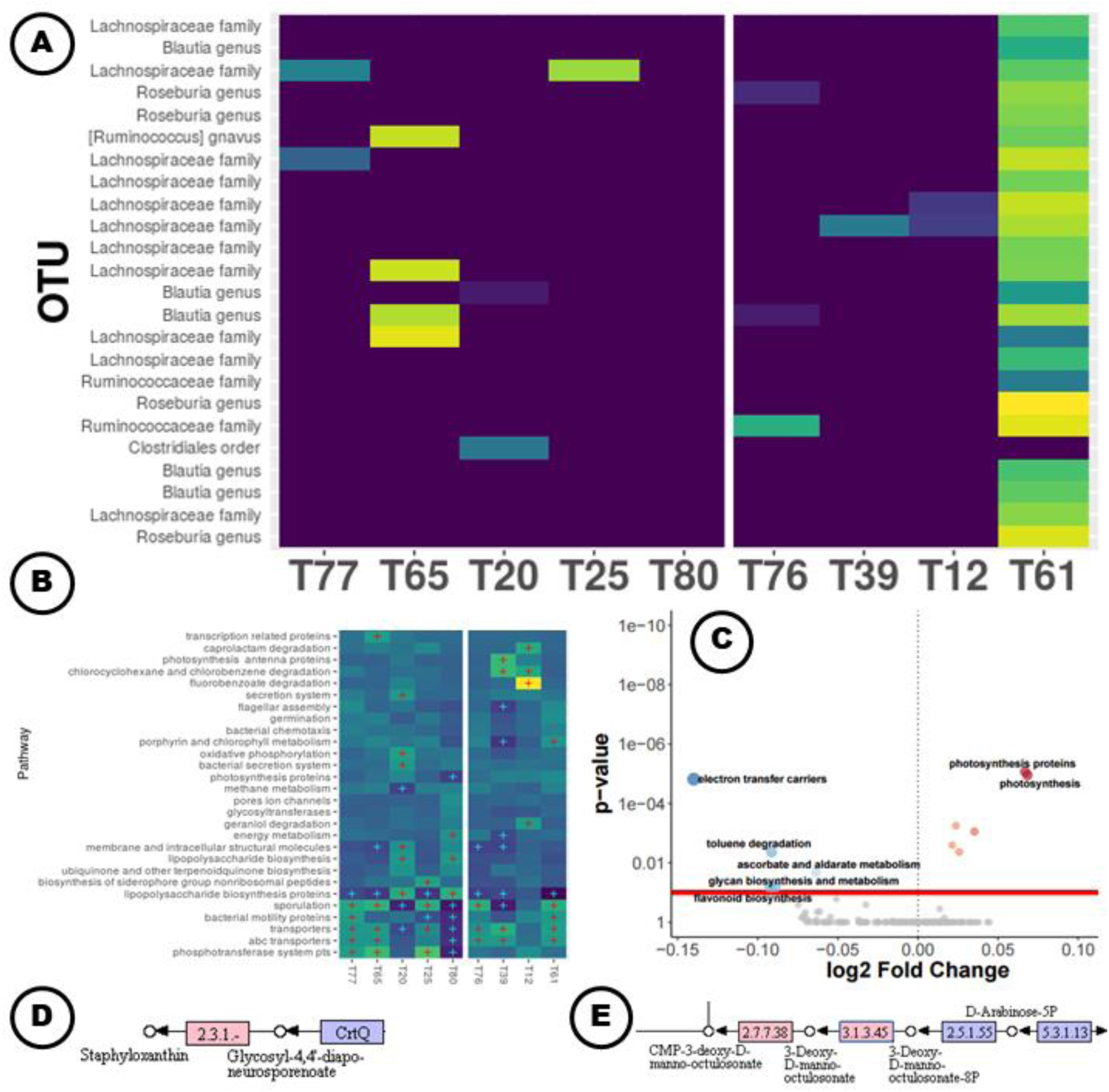
Subsection of the heatmap for AG for the topics-over-OTUs distribution (K100) in log space. Shown are the topics with 95% uncertainty intervals that do not span 0 when regressed against diet type, ordered by increasing mean regression estimate (left to right). T77 is most associated with O. T61 is most associated with V. White line signifies a shift from positive to negative mean regression estimates. Clustering was performed via Ward’s method on Bray-Curtis distances. Low probabilities (p < 1x10^-5^) are set to 0 to ease visualization. Yellow=high probability, Blue=low probability. Figure 5B. Level-3 topic-function interaction weight estimates from the multiple level negative binomial model. KEGG information was predicted via PICRUSt on the topics-over-OTUs distribution. Only the top 25 topics based on mean regression weight were chosen for the negative binomial to alleviate computational concerns. Clustering was performed via Ward’s method on Bray-Curtis distances. Red and blue crosses indicate weights for pathway-topic combinations that do not span 0 at 80% uncertainty and are positive or negative, respectively. Only pathways with at least one such combination are shown. Yellow=large positive weight estimate, Blue=large negative weight estimate. Figure 5C. Volcano plot showing DESeq2 results for differentially abundant predicted level-3 KEGG categories. Functions were predicted using PICRUSt on the copy number normalized OTU abundance table. Blue and red points represent categories significantly enriched for O and V, respectively. Gray points are categories with p-values greater than 0.1 after Bonferroni correction. Figure 5D. Glycosyl-4,4′-diaponeurosporenoate acyltransferase step (red) in carotenoid biosynthesis pathway. This gene is enriched in T77 relative to T20. Figure 5E. Lipopolysaccharide biosynthesis pathway where genes with abundances greater than 50 for T20 are colored red.

T12 contained a small yet diverse cluster of bacteria within *Acinetobacter*, a genus often associated with fermented foods and beverages [40]. Quinn *et al.* (2016), investigating the effect home-fermented foods had on human microbiota, identified enrichment of fluorobenzoate degradation pathways [41]. In our results, we found that the fluorobenzoate degradation pathway for T12 had the largest shift of any predicted pathway within a given topic (Figure 6). To further investigate the relationship between fluorobenzoate degradation pathways and diet, we performed a logistic regression (logit link) on all samples with subject ages at least 21y. Diet type and the z-scored frequency of T61 were independent variables with alcohol consumption (n_no_=837, n_yes_=3692) as the binary outcome. Both T61 (Z=3.64, β_T61_=1.10, p<0.001) and diet (Z=-6.78, β_diet_=-0.89, p<0.001) were significant, suggesting a potential relationship with fermented foods (specifically alcohol), *Acinetobacter*, and fluorobenzoate degradation.

Finally, T76 contained bacteria typically associated with a western lifestyle such as *Clostridiales* [42]. It was also enriched for *Faecalibacterium prausnitzii*, as well as butyrate production. This is significant because butyrate is not only critical in the fermentation of plant matter [43], but reduction of fecal butyrate has been implicated in obesity and a shift toward a less carbohydrate-rich diet [44]. The remaining bacteria present in this T76 cluster, *Ruminiococus* and *Roseburia*, have been shown to be elevated after fiber consumption [38].

The topics associated with the O group were enriched for LPS and secretion system pathways. A noteworthy cluster in T77 was surprisingly quite similar to the cluster in T61. Lachnospiraceae composed the majority of each cluster: 47.8% (11/23) of taxa for T61 compared to 20.6% (13/63) for T61. The functional profiles were analogous for all pathways except carotenoid biosynthesis and porphyrin and chlorophyll metabolism. A notable distinguishing characteristic is the lack of any *Roseburia* in the T77 cluster in contradistinction to T61.

Lastly, we assessed how specific parts of certain metabolic pathways were enriched or depleted within a given topic. We found that T77 was enriched for a specific gene that codes for the enzyme in the final step of the synthesis of bacterial antioxidant staphyloxanthin, a known *Staphylococcus aureus* virulence factor [45] (Figure 5D). T20 was abundant in LPS biosynthesis gene, but depleted in a subset of LPS genes key in a specific branch of the LPS pathway (Figure 5E).

### DISCUSSION

We have proposed an approach for uncovering latent thematic structure in 16S rRNA survey data that simultaneously explores taxonomic and predicted functional content. Rather than inferring functional content independently from taxonomic abundances, our approach shifts the focus to investigating within-topic functional content. Unlike other methods, by exploring our topics, we can link categories of functional content to specific clusters of taxa which can in turn be linked to sample features of interest. For example, like Gevers *et al.*, we detected a relationship between membrane transport genes and CD+, but our approach also allowed us to determine which bacteria (OTUs belonging to Enterobacteriaceae) were the prime contributors to the enrichment of membrane transport genes. Moreover, the pathogenic set of bacteria reported by Gevers *et al.* (*Haemophilus* spp., *Neisseria*, and *Fusobacteria*) contributed less to the abundance of membrane transport genes. By applying statistical approaches on the dataset as a whole, as is typical, the apparent relationship between membrane transport genes and specific clusters of bacteria would be lost.

We have also shown that our approach drastically reduces the dimensionality of two high-dimensional sources of information, taxonomic abundances and functional content, increasing the ease in which these data can be interpreted. For instance, we can identify gene sets of interest from noteworthy topics. For the Gevers *et al.* dataset, we determined that T15 is (1) associated with CD+ samples; (2) dominated by a cluster of bacteria previously associated with CD; and (3) uniquely enriched for a subset of LPS synthesis genes. With a gene profile from a topic of interest, one could focus on gene subsets associated with topic-specific bacterial clusters that are known disease biomarkers, which in turn may facilitate targeted approaches for future research endeavors.

Lastly, our complete pipeline is computationally manageable. Fitting the STM to nearly 5000 samples from the AG dataset reaches convergence in minutes. Functional prediction via PICRUSt also only takes minutes (using our C++ implementation in themetagenomics). Inferring topic-function interaction effects via our multilevel, negative binomial regression approach is comparatively slower, however, taking hours for large datasets (*e.g.,* AG). This is because we implement this model in the probabilistic programming language Stan, which uses Hamiltonian Monte Carlo. Maximum likelihood (a much faster alternative) generally fails to converge for these data, although the regression weight estimates tend to be quite similar based on our experience.

We present our approach at a time when novel means to analyze complex microbiome abundance data is called for. Current methods often link the abundance of a single OTU across samples to a sample feature of interest. These methods routinely identify important subsets of taxa, but ignore OTU co-occurrence. Network methods overcome this concern, but instead fail to do so while including sample information within the model. Consequently, they are incapable of directly linking sections of the OTU correlation network with sample features of interest. Constrained ordination methods, such as canonical correspondence analysis, do in fact couple inter-community distance with sample information, but the user is limited to specific distance metrics (*e.g.,* Chi-squared) and must follow key assumptions (*e.g.,* the distributions of taxa along environmental gradients are unimodal) [46]. Moreover, interpretation of biplots becomes increasingly difficult as more covariates are included, and, unlike our approach, linking key groups of taxa with functional content is not straightforward.

The ability to make meaningful inferences is further limited by the fact that microbiome data is often inadequately sampled (thus justifying some type of normalization procedure), compositional (due to normalization), sparse, and overdispersed. Compositional data restricts the appropriateness of many statistical methods due to the sum constraint placed across samples. SPIEC-EASI provides a robust network approach for overcoming compositional artifacts in an attempt to infer community level interactions. We compared our within-topic taxonomic profiles to the first and second order connections identified by SPIEC-EASI, to which we found coherence between the two approaches, suggesting a topic model approach for compositional data is in fact appropriate.

Others have explored the use of Dirichlet-Multinomial models, which are well equipped at managing overdispersed count data [47–49]. The fact that Dirichlet-Multinomial conjugacy is exploited for the topics-over-OTUs component of many topics models reflects their suitability for abundance data. We selected the recently developed STM for our workflow because of its ability to not only utilize sample data prior information in the flavor of the Dirichlet-Multinomial regression topic model, but also its ability to capture topic correlation structure and apply partial pooling over samples or regularization across regression weights.

Because normalization is also a chief concern when analyzing sequencing abundance data [27], we found it imperative to determine a suitable approach. In the original LDA paper, the generative process assumed a fixed document length *N*, but *N* was considered a simplification and could easily be removed because it is independent of all other components of the model. This allows for the possibility of more realistic document size distributions [15]. Giving this fact, in a companion paper, we performed simulations to determine a suitable normalization strategy (Woloszynek *et al.*, *in prep*). Our simulations suggested that predefined subcommunities concentrate with high probability to extracted topics and that no library size normalization was required to maximize power or the ability to infer taxonomic structure. Thus, a topic model approach provides a direct, suitable procedure for inferring the subcommunity configuration. The variance stabilization through DESeq2, while potentially ideal for large sample sizes with adequate signal, seemed to dampen the ability to identify topic-sample associations. Despite performing well at mapping subcommunities to topics, the rarefied approach suffered from reduced power when identifying topics with large covariate effects. Taken together, we concluded that raw abundance data could be adequately modeled in our approach.

There are limitations to our approach. First, the topic-function inference step currently scales poorly in terms of computation time for large numbers of topics, which may be more important as datasets continue to grow in size. Regularization and sparsity-inducing priors help limit the number of important topics; hence, exploring only a subset of topics during the final regression step can offer substantial speed improvements at little cost, but utilizing the complete set of topic information would be ideal. Also, we used Hamiltonian Monte Carlo via Stan. As of now, Stan lacks within-chain parallelization, although that may change in upcoming releases (see Stan MPI prototype). Alternatively, other posterior inference procedures such as variational inference using software packages such as Edward may provide additional speed enhancements [50]. Second, we are capable of separately estimating the uncertainty in our topic model, the hierarchical regression model, and the functional predictions from PICRUSt, but we currently do not propagate the uncertainty throughout the pipeline. Doing so would improve downstream interpretation with better estimation of the topic-sample covariates and pathway-topic effects, which in turn would greatly improve one’s confidence with using within-topic gene sets. Third, we do not incorporate phylogenetic branch length information, which could lead to more meaningful topics.

## LIST OF ABBREVIATIONS

AG: American gut
BMI: body mass index
CD: Crohn’s disease
IBD: inflammatory bowel disease
LDA: latent Dirichlet allocation
LFC: log-fold change
LN: logistic Normal
LPS: lipopolysaccharide
OTU: operational taxonomic unit
PCDAI: pediatric Crohn’s disease activity index
PPD: posterior predictive distribution STM, structural topic model
STM: structural topic model

## AVAILABILITY OF DATA AND MATERIAL

The datasets generated and/or analyzed during the current study are available in the tm_pipline github repository, https://github.com/sw1/tm_pipeline

## AUTHORS’ CONTRIBUTIONS

SW chose and preprocessed the datasets; developed the pipeline, modeling approach, and statistical tests; designed the topic-function NBR model; and fit all models and performed all tests. JM and SW conceived of the topic-function prediction approach. SW and GS designed simulations for normalization and determined the approach to assess posterior uncertainty. SW, JM, GR, and MO assisted in interpretation and identifying ways to further validate the pipeline. GR supervised the project. All authors contributed to the writing of the manuscript. All authors read and approved the final manuscript.

## ACKNOWLEDGEMENTS

We would like to thank Zhengqiao Zhao for his feedback on the manuscript and help in developing the simulation. We would also like to thank the Casey Greene Lab for their information on continuous integration and reproducibility, which was invaluable for developing themetagenomics.

